# Live analysis and reconstruction of single-particle cryo-electron microscopy data with CryoFLARE

**DOI:** 10.1101/861740

**Authors:** Andreas D. Schenk, Simone Cavadini, Nicolas H. Thomä, Christel Genoud

## Abstract

Efficient, reproducible and accountable single-particle cryo-electron microscopy structure determination is tedious and often impeded by lack of a standardized procedure for data analysis and processing. To address this issue, we have developed the <u>F</u>MI <u>L</u>ive <u>A</u>nalysis and <u>R</u>econstruction <u>E</u>ngine (CryoFLARE). CryoFLARE is a modular open-source platform offering easy integration of new processing algorithms developed by the cryo-EM community. It provides a user-friendly interface that allows fast setup of standardized workflows, serving the need of pharmaceutical industry and academia alike who need to optimize throughput of their microscope. To consistently document how data is processed, CryoFLARE contains an integrated reporting facility to create reports.

Live analysis and processing parallel to data acquisition are used to monitor and optimize data quality. Problems at the level of the sample preparation (heterogeneity, ice thickness, sparse particles, areas selected for acquisition, etc.) or misalignments of the microscope optics can quickly be detected and rectified before data collection is continued. Interfacing with automated data collection software for retrieval of acquisition metadata reduces user input needed for analysis, and with it minimizes potential sources of errors and workflow inconsistencies. Local and remote export support in Relion-compatible job and data format allows seamless integration into the refinement process. The support for non-linear workflows and fine-grained scheduling for mixed workflows with separate CPU and GPU based calculation steps ensures optimal use of processing hardware. CryoFLARE’s flexibility allows it to be used for all types of image acquisitions, ranging from sample screening to high-resolution data collection, and offers a new alternative for setting up image processing workflows. It can be used without modifications of the hardware/software delivered by the microscope supplier. As it is running on a server in parallel to the hardware used for acquisition, it can easily be set up for remote display connections and fast control of the acquisition status.

## Introduction

The advent of direct electron detectors ^1–3^ in combination with data acquisition automation (e.g. with EPU, SerialEM ^4^, Leginon ^5^) and new algorithmic advances in image processing have contributed to the establishment of single-particle cryo-EM as a successful and attractive tool for structure determination of proteins and protein complexes ^6,7^. To obtain high resolution cryo-EM structures, data in the order of terabytes are routinely collected, analyzed and processed. The large amount of data produced is a growing challenge as it must be shared, stored, transferred, viewed and archived. To overcome this challenge automated tools for quality assessment, data processing, data handling and reporting are required. In parallel, it is necessary to keep hardware and software updated in order to follow the fast developments in the single-particle cryo-EM field. Therefore, a data processing workflow needs to be fully open and easy to modify in order to integrate newly developed algorithms at the pace they are offered to the community.

In this context we introduce CryoFLARE to perform live analysis and reconstruction of data in parallel to data acquisition. It is an open-source software written in C++/Qt5. It shares similarities with processing frameworks as e.g. Focus ^8^ or Warp ^9^ but focuses more on providing a standardized and yet flexible graphical workflow with strong integration of data acquisition and reporting suitable to document the data integrity prior to publication. CryoFLARE does not require programming knowledge for setup and use. It provides an intuitive user interface displaying multiple parameters during acquisition such as ice thickness, devitrification level and number of particles picked. Acquisition parameters as e.g. camera type, nominal defocus, phase plate information and exposure time are automatically imported from the acquisition software without user input, ensuring coherence and accuracy of acquisition metadata.

CryoFLARE is agnostic to the actual algorithms used for processing. Instead it provides an interface to communicate with existing processing scripts or programs by transferring parameters through standard input and output channels.

CryoFLARE allows flexible integration of newly available algorithms instead of depending on a fixed pipeline or implementation. Its real-time visualization capabilities allow live monitoring of data acquisition and can serve as a tool for ensuring microscope performance and data quality. CryoFLARE provides a fast and rigid testing ground for evaluation and comparison of new processing strategies or algorithms. It supports optimization and acquisition with Volta phase plates (VPP) by monitoring and controlling the positions used on the VPP and tracking its performance

Users can assess the quality of their data and adapt the image acquisitions conditions quickly at the beginning of their imaging sessions. Micrographs to be kept and re-processed can be selected interactively. CryoFLARE can also transfer the data to the image analysis software of choice easily. For the experts in image analysis responsible for the data acquired in a laboratory, the clear and documented structure of CryoFLARE allows a fast adaptation of the workflow to special processing needs, e.g. switching to different CTF determination software for tilted data collection, or automatic phase shift fitting if a phase plate is inserted during acquisition. The graphical user interface makes users aware of the micrograph quality and data acquisition optimization as the feedback is instantaneous and displayed graphically. Its intuitive design allows going through micrographs and adapting particle picking strategies while data is acquired. The detailed aspects are illustrated and described below.

## User interface

The graphical user interface (Figure 1) is organized into two separate pages. The first page is dedicated to visualizing micrograph related results, while the second page is used to aggregate information on the grid square level.

**Figure 1:**
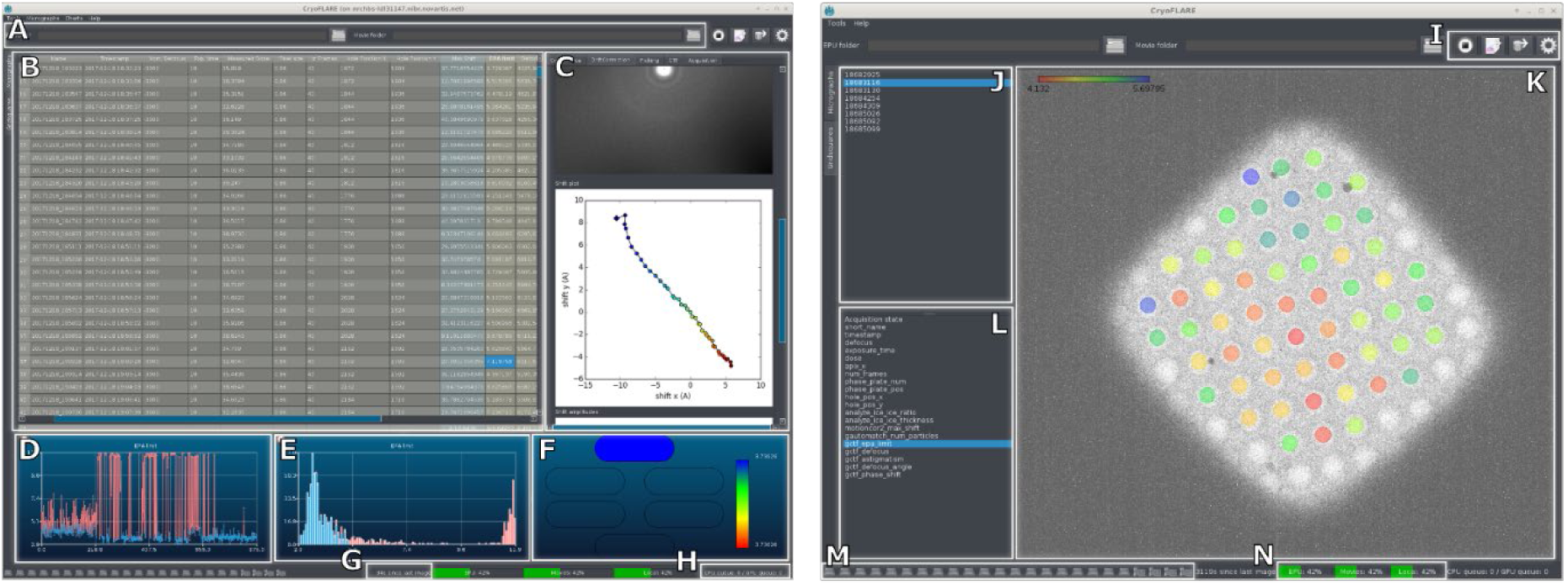
Graphical user interface: (A) Project and movie folder selection. (B) List of micrographs displaying results from acquisition and processing. (C) Detail panel showing results from currently active micrograph as well as input form for user defined parameters. (D) Linear plot. (E) Histogram. (F) Phase plate heat plot. (G) Counter indicating time since last acquisition. (H) CPU and GPU queue (I) Buttons for export, reporting, start/stop and settings. (J) Grid square selection. (K) Grid square display with heat plot overlay. (L) Parameter selection for heat plot. (M) CPU and GPU activity display. (N) Disk usage display.

The micrograph page consists of multiple areas: The table in the top left (Figure 1B) lists each acquired micrograph and displays the corresponding metadata from the acquisition software (e.g. pixel size, dose, etc.) live as well as metadata created by processing tasks (e.g. defocus, drift, etc.). The design of the table was inspired by the IPLT diffraction processing pipeline ^10–12^. The table also serves as a tool for selecting individual micrographs or micrograph groups for export. The panel on the right-hand side (Figure 1C) shows details about the micrograph currently selected in the micrograph table. The results are grouped by processing task (e.g. motion correction, particle picking, etc.) and consist of metadata or images and plots associated to the task. Each task is displayed in a panel and all panels are organized as tabs that can be selected. In addition, the panel allows modifying the input parameters for the displayed task. For example, in the particle picking panel, the picking algorithm and box size can be modified at any time during processing. The bottom panels (Figure 1D-F) provide live graphs calculated over the whole data set. From left to right the first graph shows a linear plot, following the progress of the data acquisition with the X-axis representing the micrograph number. The parameter to be plotted in Y can be chosen by selecting a parameter column in the micrograph table (for example astigmatism). The second graph shows a histogram distribution of the same selected parameter. The third panel shows a heat plot and allows mapping the selected parameters onto individual VPPs and VPP positions, if the dataset was acquired with a VPP present. The graphs can be freely zoomed and are updated in real time with each processed micrograph. In addition, graphs can be used for interactive selection of micrographs by the user (see Data selection below). Selected micrographs are plotted in blue, while deselected micrographs are plotted in red. In addition to the standard plots, two freely selectable parameters can also be plotted against each other in a scatter plot to evaluate their correlation. For example, we found it useful to plot ice thickness versus the maximum resolution of the CTF fit, as thinner ice often correlates with better data quality(Figure 4C).

**Figure 2:**
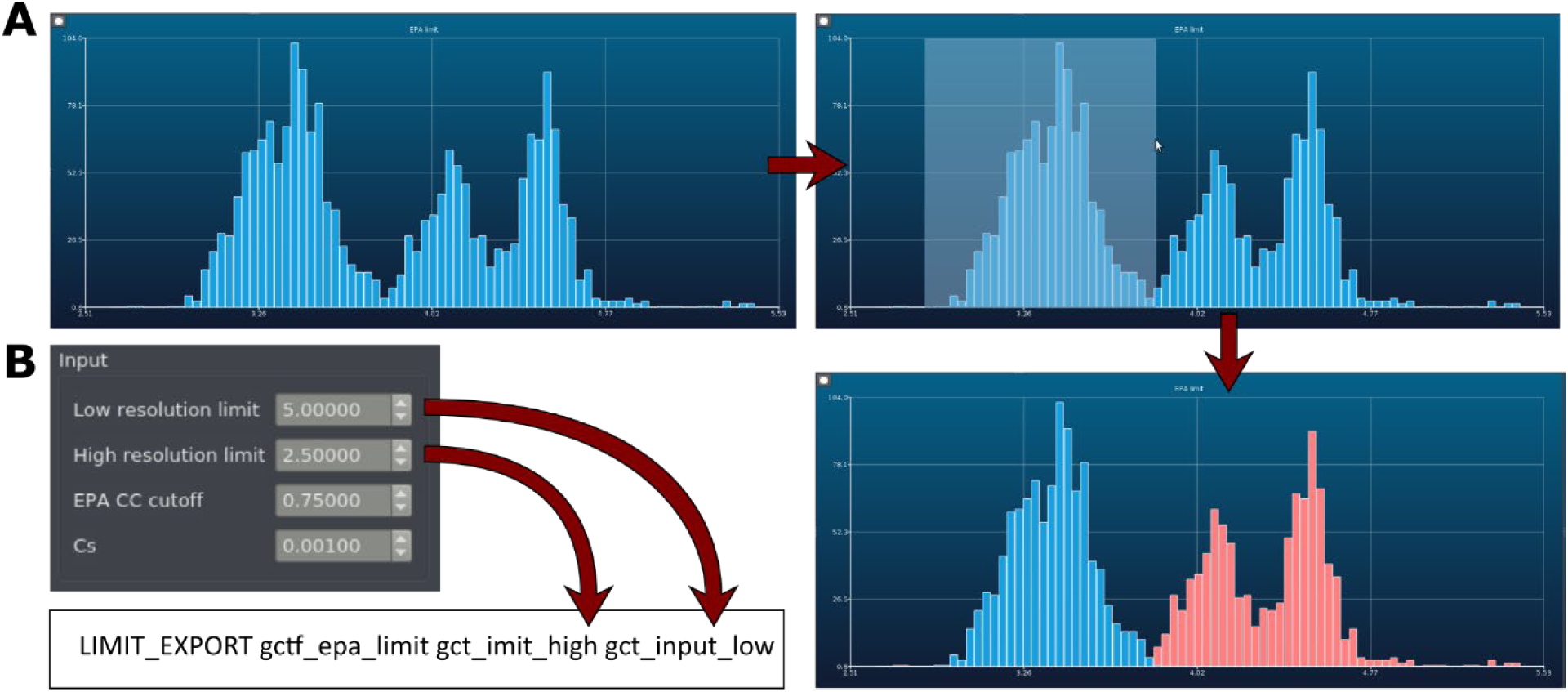
Data selection: (A) Data can be interactively selected with the mouse on the real time graphs. (B) Data can be selected based on user defined limits, which are then used for selection within the processing script. The shown example selects all images that show a GCTF EPA limit (stored in the bash variable gctf_epa_limit) between 2.5 Å and 5.0 Å for export.

**Figure 3:**
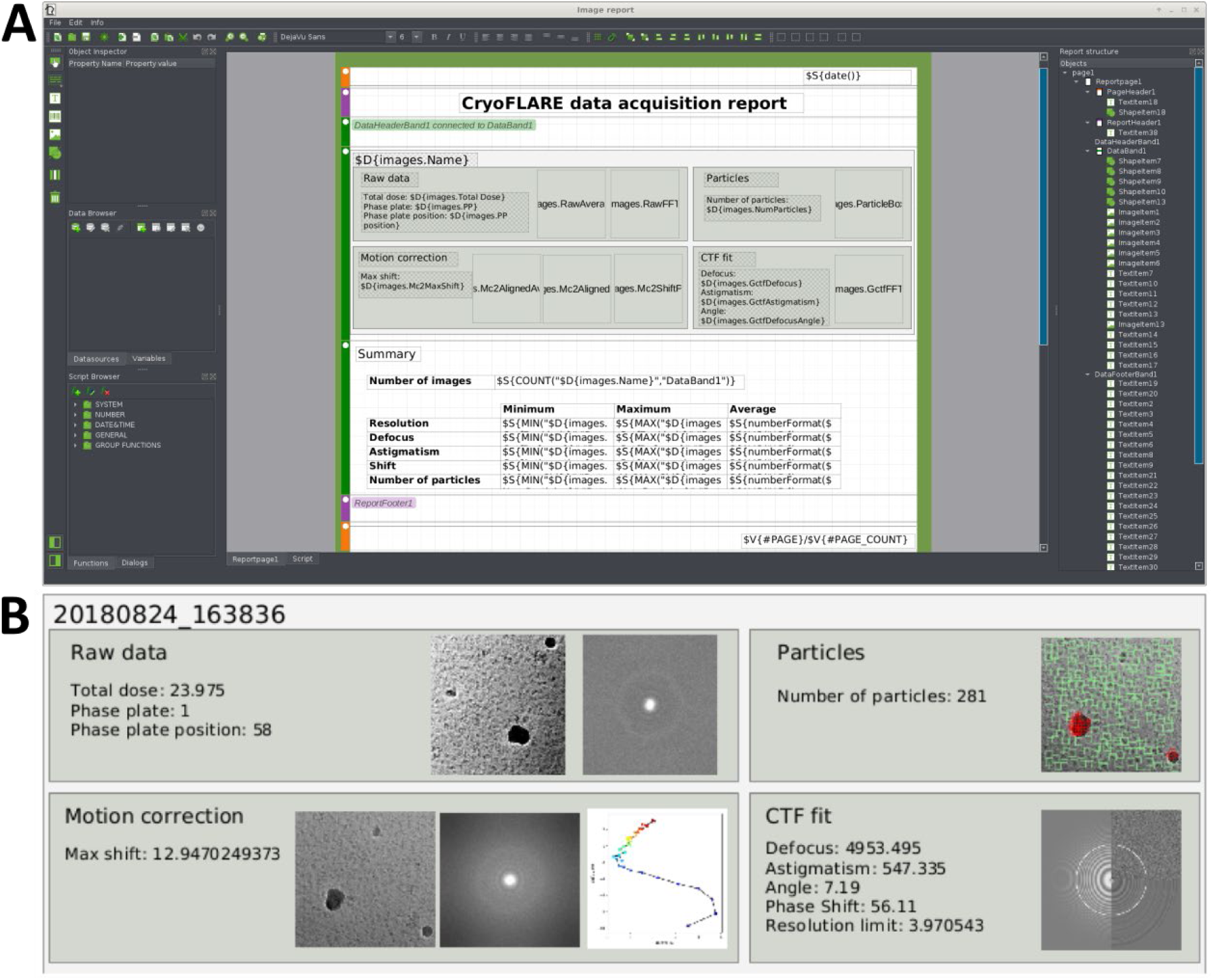
(A) Report builder. (B) Sample report output for single micrograph.

**Figure 4:**
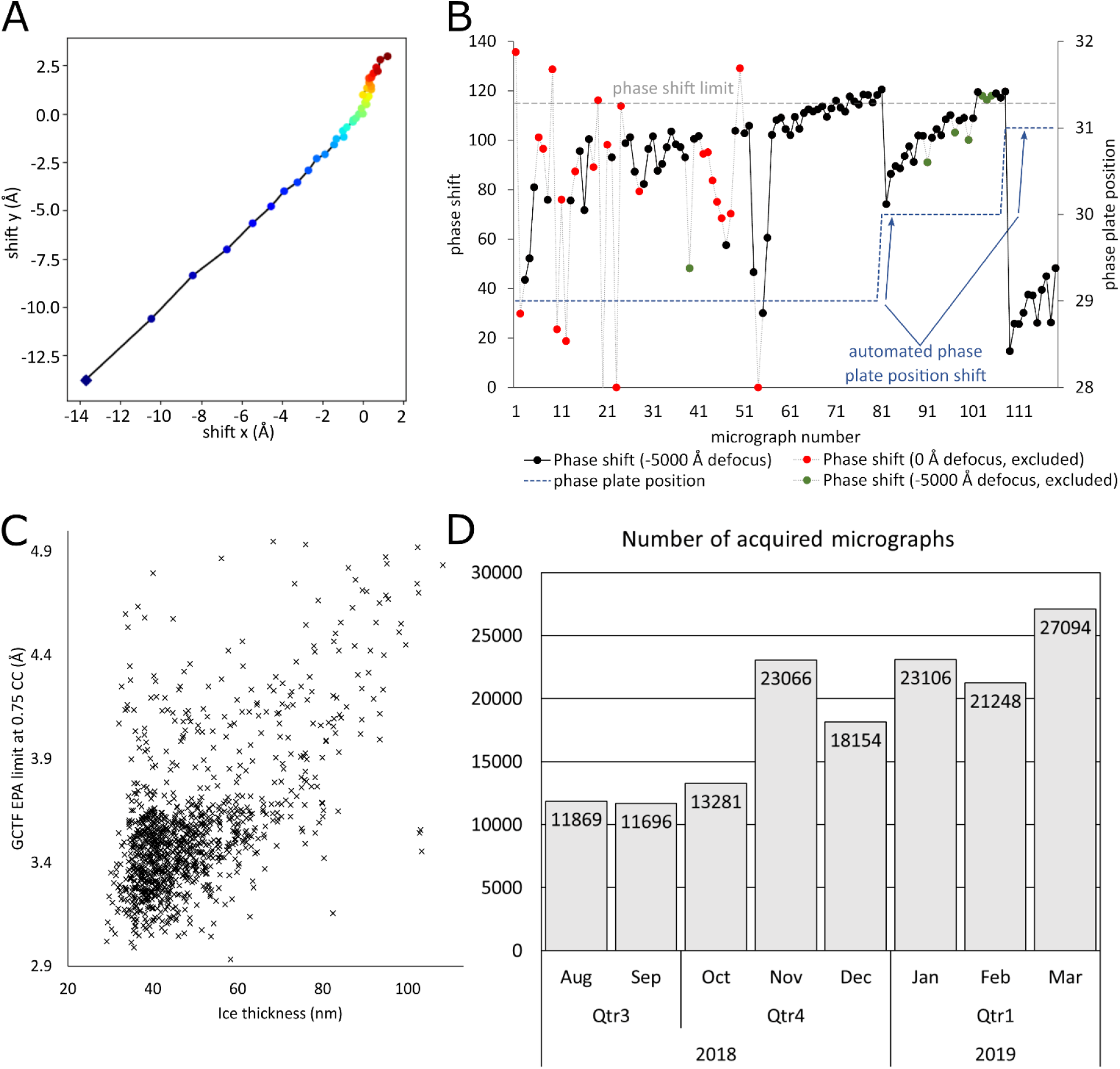
(A) XY plot of frame shifts during motion correction with a blue diamond marking the first frame. (B) Phase shift of individual micrographs. Micrographs showing problems during CTF fitting are automatically excluded. Most micrographs were excluded due to low contrast because of in focus acquisition (red), whereas few micrographs showed other fitting problems (green). The phase shift values were uses to automatically control phase plate movement. The phase plate position was shifted at a median phase shift above 115° within 10 micrographs. (C) Scatter plot correlating ice thickness to GCTF EPA limit. (D) Number of acquired micrographs over a time span of 8 months.

The grid square page shows an overview of the recorded grid squares. The overview micrograph can be overlaid with a heat plot, mapping any of the results onto their relative foil hole position within the grid square.

The processing of each micrograph is split into separate tasks for each step of the processing (drift correction, CTF determination, etc.). The processing tasks to be performed on each micrograph can be defined in the settings dialog box. The tasks are organized into a tree instead of a linear workflow to allow parallel execution of independent processing tasks. For each task a set of input and output parameters can be defined to facilitate data exchange with the processing script or program.

## Features

### Import

CryoFLARE supports three separate import options to handle differences in folder organization by different data acquisition packages. Data can either be organized in 1) an EPU folder structure, with full support for multiple Images-Discs and grid squares, 2) a flat folder structure providing micrographs and EPU metadata xml files in a single folder, allowing reimport of already processed EPU data, or 3) a flat folder structure with optional metadata in json format, allowing processing of data from 3^rd^ party data acquisition packages as e.g. SerialEM^4^ or Leginon^13^. Extracting and using metadata directly from the data acquisition software allows users to concentrate on data analysis as it avoids the burden of manually entering already determined parameters again into the analysis software. This automation reduces the source of possible errors by misentering data. For EPU data, CryoFLARE reads the metadata associated to grid squares foil holes as well to allow data analysis on the grid square level.

CryoFLARE supports file creation and modification detection on remote shares (e.g. CIFS or NFS), hence enabling direct import from a remote location as for example a microscope or camera PC. Support for remote data locations allows to separate the microscope network segment (in general accessible to external parties for remote support) from the internal network environment which might contain confidential data (Figure 6B). Fencing off the data collection hardware reduces the attack surface on the internal network and lowers the risk of exposure of internal data.

**Figure 5:**
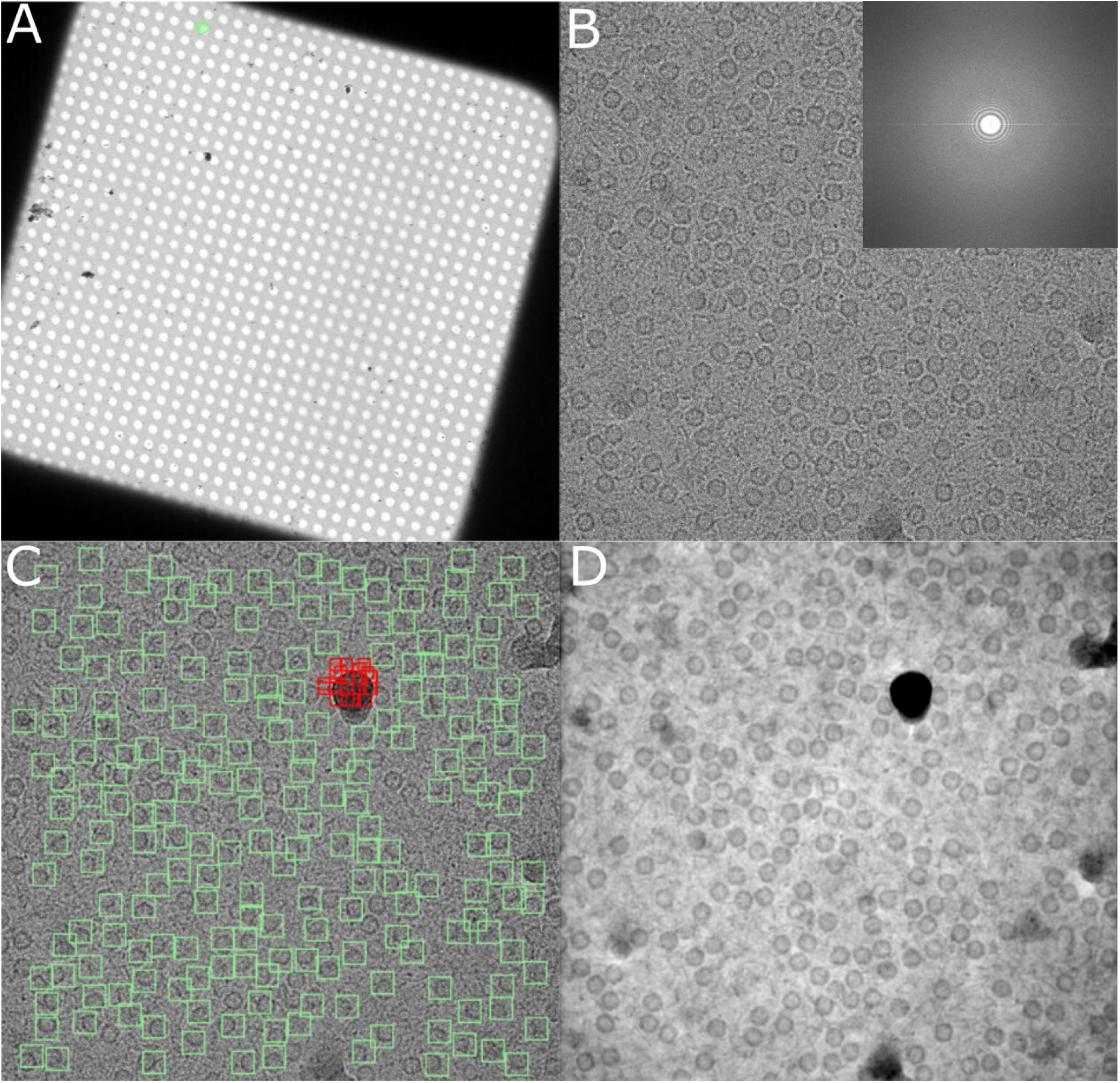
Sample workflow: (A) Foil hole position of recorded micrograph indicated in green on grid square. (B) Motion corrected micrograph with FFT in inset. (C) Green boxes indicating picked particles. Red boxes indicating excluded areas. (D) Deconvoluted micrograph.

**Figure 6:**
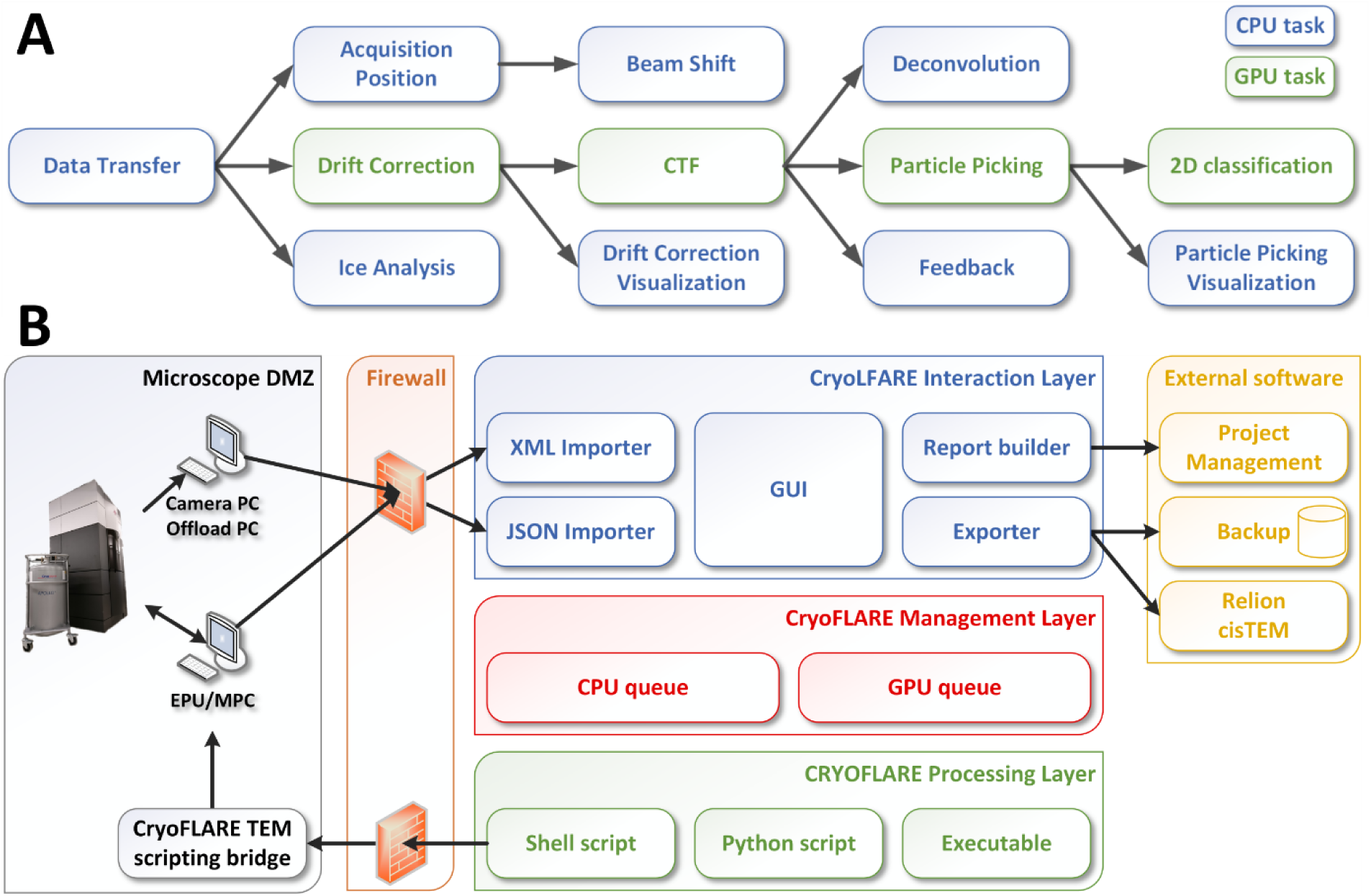
Implementation: (A) process tree. (B) Software layers.

### Data selection

To provide an optimal data set for further processing it is invaluable to have a selection regime in place to remove bad micrographs during live processing. CryoFLARE supports the user by providing three easy to use selection modes. 1) The list of micrographs can be sorted based on any of the results and then micrographs can either be selected individually or all micrographs above/below a certain limit can be selected or deselected. 2) Subsets of micrographs can be interactively selected (Figure 2A) in a live plot. 3) Micrographs can automatically be selected or deselected based on an upper and lower limit for a parameter within a processing script (Figure 2B, Supplementary Figure 2A). The ability to select a subset of good data for subsequent export leads to faster data transfer, reduced storage requirements and facilitates classification and refinement, therefore supporting a sustainable high quality and high throughput processing workflow.

### Reporting

For a single-particle EM structure determination pipeline, especially if it is run in a facility setup, it is becoming increasingly important to provide adequate documentation on data acquisitions and processing steps that are performed. The documentation provides accountability and project continuity by simplifying handover of data between multiple people involved in a project. CryoFLARE supports this effort by providing a reporting facility that can be tailored to individual use cases. Report templates can be assembled in a graphical user interface (Figure 3A), where parameters or images from data acquisition or processing can be selected interactively. At the end of the data acquisition a report in pdf format (Figure 3B) can be created using the template assembled beforehand.

In addition, the reported values can also be saved as a list of comma-separated values (CSV) or in json format. The saved files allow creation of graphs for publication in external software, as illustrated in the example plots in Figure 4B and C.

### Processing

For each acquired micrograph a set of processing tasks is performed. The metadata generated during acquisition and processing is carried over to subsequent tasks, making it easy to set up processing pipelines with minimal user interaction. The metadata together with the processing settings are stored permanently within the processing folder to ensure persistence of the data across multiple program executions.

The CryoFLARE program itself does not include any processing algorithms or user interface elements fixed to a certain metadata parameter. Instead it provides an API (see Scripting interface) to dynamically execute user defined processing tasks. This design was chosen to allow users the flexibility to define their own workflows and to enable them to include newly developed processing algorithms, instead of forcing users to use one specific implementation. The workflow is saved and loaded together with the rest of the settings. CryoFLARE saves the settings within the processing directory and opening CryoFLARE again in the same directory at a later stage will load them again to restore the workflow. In addition, settings can be saved as account wide default, which are then used whenever CryoFLARE is started in a processing directory not containing any settings.

Using this generic interface, we set up a processing workflow which is distributed together with CryoFLARE, containing the following advanced features:

#### Multi-step CTF determination

The CTF determination script uses Gctf ^14^ to determine the CTF parameters in an in-house developed two-step process. In a first step a fit is performed with a defocus step size of 500 Å. The search range is dependent on the nominal defocus, as provided by the data acquisition software. A lower limit of half the nominal defocus and a upper limit of 2.5 times the nominal defocus is used. In a second step, the CTF parameters are refined using a defocus step size of 100 Å within a range of 0.9x to 1.1x of the average defocus determined in step one. For equiphase averaging resolution estimation of the final fit, a user defined cross correlation cutoff is can be set. A default cutoff value of 0.75 is used. The resulting CTF parameters are stored into a star file for further use in 2D classification in Relion^15^ or Cryosparc^16^. The CTF parameters are also used to deconvolute the micrographs (Figure 5D) using a script implemented in IPLT ^10,11^ following the algorithm described in Tegunov & Cramer^9^.

#### Particle picking

Particles are picked on each micrograph using Gautomatch (https://www.mrc-lmb.cam.ac.uk/kzhang/). The Gautomatch output is parsed and particle positions are indicated on the micrograph in green while rejected positions are indicated in red (Figure 5C).

#### Ice thickness determination

The ice thickness of each acquired micrograph is automatically estimated using the aperture limited scattering methodology laid out in Rice et al.^17^. The python script performing the calculation uses EMAN2 ^18^ to extract the relevant micrograph data. The sole input that must be provided by the user is a reference micrograph without a sample recorded with identical illumination as used for imaging.

#### Devitrified ice detection

Devitrification is detected using a homemade script by calculating the ratio between the signal intensity in a micrograph at 3.89 Å and 5.0 Å. Deviation of the ratio from 1.0 can signal possible devitrification of the sample. The python script uses EMAN2^18^ to calculate the rotational average of the Fourier transform of the micrograph. From the rotational average an ice band at 3.89±0.2 Å and a reference band at 5.0±0.2 Å are extracted. The maximum value in the ice band (*V*_*ice*_) and the reference band (*V*_*ref*_) are determined and the ratio 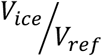 is calculated. A ratio significantly over 1.0 reproducibly indicates micrographs with devitrified ice.

#### Amplitude graphs and 2D plots for motion correction

The script provided with CryoFLARE for motion correction uses MotionCor2 ^19^. To visualize the correction 2D scatter plots depicting the frame to frame shift vectors (Figure 4A) and amplitude plots depicting the absolute amount of shift between frames are created using matplotlib ^20^.

#### Tracking of acquisition position

CryoFLARE reads the EPU metadata files for each acquired micrograph to extract the actual position of the acquisition in reference to the grid square. The position is drawn as a green circle on the grid square overview image.

#### Aberration present image shifting support

CryoFLARE creates a 2D vector diagram with matplotlib, indicating the applied image shift for each acquisition. This information can be used to identify individual acquisitions in EPU template setups used for aberration present image shifting^21^ (APIS). In addition, CryoFLARE tracks and writes template position ID into the star file containing the CTF information for each micrograph, therefore allowing grouping by template position for beam tilt refinement (e.g. in Relion).

#### Data aggregation

To monitor performance of the instrument over time, each exposure is logged in csv format in a file. For each exposure the sample position, phase plate position (if a phase plate was used), maximum drift, ice thickness and quality, CTF parameters including phase shift (if applicable) and number of detected particles are stored. The csv data can be used for plot generation as for example shown in Figure 4D.

#### 2D classification

Automatic 2D classification can be triggered once a user-defined number of particles are picked. The 2D classification will then be carried out on particles from selected micrographs (see Data selection) by filtering out deselected micrographs from all star files involved. The particle coordinates determined during particle picking are used to extract particle boxes with relion_preprocess. The extracted particles are then classified in 2D using relion_refine. The results are displayed as a montage created by Sparx/EMAN2.

### Feedback control

CryoFLARE provides a Windows service to enable access to Advanced TEM Scripting from a remote computer, where CryoFLARE is running. This is used in our setup to automate changing of the phase plate position. A script checks the CTF fits of a user defined number (10 in this example) of the most recent micrographs, and once the median phase shift exceeds a user defined threshold (115° in this example), the phase plate is shifted to the next position (Figure 4B). Automation of phase plate movement ensures that micrographs are recorded in an optimal phase shift range to maximize micrograph contrast.

### Export

The export of the data is split into two separate elements: 1) the file and destination selection and 2) the data filtering and actual transfer. The file destination selection provides the user with a dialog, where he can select which files to export for the chosen micrographs. The files are split into three separate lists for raw data, processing data, and shared files (e.g. star files listing all micrographs). The user can also browse for a local or remote destination folder as well as a separate destination folder for the raw data (e.g. for separate backup). Any folder reachable by ssh can be browsed and selected as destination.

Once the data transfer is started, the exporter will generate a list of micrographs with associated processing data to be transferred. Only data from selected micrographs that are fully processed and selected by the user, or the processing script, will be transferred. As a second step the exporter parses all star files containing data from multiple micrographs and filters out micrographs that are not exported. The filtering step is an important part of the workflow to ensure only useful data is exported and thereby reducing transfer times and storage requirements.

The selected data is exported to the destination using several threads in parallel to optimize data transfer speed.

## Sample workflow

Here we present a sample workflow in CryoFLARE as it is currently used in our facility. An overview of the workflow can be seen in Figure 6A. The first task is to transfer the averaged micrograph and metadata from the microscope PC and the raw stack and gain refence from the Gatan PC (for a K2 camera) or Offload PC (for a Falcon III camera) to the processing server. Once the transfer is complete, as next steps motion correction, ice analysis and visualization of the acquisition position followed by image shift visualization are performed in parallel. Visualization of inter-frame motion is performed in a script separate from the actual motion correction to avoid blocking GPU resources longer than necessary. The visualization is run in parallel with the next GPU task, which is the CTF determination. Upon completion, a feedback task is started allowing automatic skipping to the next phase plate position once a phase shift threshold has been reached. In parallel, processing is continued with GPU accelerated particle picking. Once particle picking is finished the visualization of particle positions is performed and at the same time 2D classification is started once enough particles were picked.

## Implementation

CryoFLARE is organized into three different layers (Figure 6B). The interaction layer handles the interaction with the user, the acquisition software, backup locations and further downstream processing like classification and refinement. The management layer is responsible for scheduling the processing and optimizing the use of the computing infrastructure. The processing layer oversees running the processing tasks and interfacing with the external processing scripts and programs.

### Scheduler

At the core of the CryoFLARE implementation sits the scheduler. The scheduler contains two stacks to separate scheduling of GPU and CPU tasks. The tasks are processed in last-in-first-out (LIFO) order to give highest priority to the most recent data. The number of processing slots assigned for each stack can be configured separately, as on most systems the number of CPU cores does not match the number of GPUs. This setup allows to fully use the CPU resources without forcing multiple tasks to share the same GPU, given that this could lead to instabilities in the processing (e.g. with MotionCor2 version 1.0.1 and above). Breaking up the micrograph processing into separate tasks allows to define non-linear workflows and with it parallelization within a micrograph processing workflow (Figure 6A). In addition, the task-based scheduler allows allocation of GPU resources to be limited to tasks utilizing it, instead of allocating the resources for the full workflow. Tasks are spawned as separate processes using the QProcess interface.

The scheduler checks the completion of each tasks and tracks that information by setting flags in the metadata object (see below) associated with a micrograph.

The organization of the scheduling into self-contained tasks together with the fact that CryoFLARE also reads in metadata for grid squares would also allow to run tasks on the grid square level in the future to e.g. predict good sample areas before data acquisition and transfer back that information into the data acquisition software.

### Metadata

Metadata is internally stored as Json object. The metadata associated with each micrograph consists of data acquisition parameters, result parameters generated during data processing within CryoFLARE, file paths for files generated during processing as well as completion states of the processing tasks. Upon each change of metadata, the update is written out to a binary encoded Json file in order to ensure data persistence. Upon restart of CryoFLARE in a pre-existing processing directory the binary encoded metadata is read in to restore the processing state.

### Scripting interface

A scripting interface is provided within CryoFLARE to transfer metadata between user interface and processing scripts or programs. Upon start of a processing task by the scheduler all existing metadata items for the micrograph to be processed are transferred to the processing script/program as key=value pairs on the standard input. For python and bash script an adapter is provided that automatically translates the input into script variables named after the metadata keys (Supplementary Figure 1A). The user interface settings allow definition of additional metadata items to serve as user defined input for processing tasks.

Output from the processing is transferred back to CryoFLARE in a similar way on the standard output. In addition to output parameters, files can be defined as well. Three types of files are distinguished: output files, raw data files, and files shared between all micrographs (e.g. micrographs star files). The distinction enables more selective data export (see Export). For bash and python scripts convenience functions are provided to easily define output parameters (Supplementary Figure 2C) and files (Supplementary Figure 1B,C). It is important to note that any output metadata is automatically made available as input to subsequently running child tasks.

To facilitate interaction with Relion for subsequent processing, a set of scripting functions is provided to create Relion job folders and pipeline files as well as creating and adding data to star files (Supplementary Figure 2B). The scripting functions take care of file locking in order to make them safe to use during parallel processing of multiple micrographs.

### Feedback API

To transfer back information and control the microscope CryoFLARE provides a bridge to Advanced TEM scripting. The bridge consists of a Windows service implemented in Python using the pywin32 interface. It takes commands from a configurable control file, which are then translated and passed on to the Advanced TEM scripting COM interface, which is accessed through the comtypes module. Use of a file-based control connection allows to easily pass firewalls. CryoFLARE processing tasks can write commands to the control file (located on a mounted remote share) to control microscope function and therefore feedback processing information to improve data collection.

### Importer

The data import facility consists of two separate elements: a file and folder listener to watch for newly acquired data and a metadata reader to retrieve metadata information from the data acquisition software. Both elements are encapsulated in an importer class to allow easy adaptation to different folder and metadata structures.

The file and folder listener is a drop-in replacement for QFilesystemWatcher, reimplemented from scratch using a polling scheme to support listening on remote shares that do not provide full file change notification support. The listener is started in a separate thread to avoid unnecessarily blocking the user interface and supports watching multiple folder within a directory structure in order to fully observe nested directories as created by EPU.

Two metadata readers are implemented, one for XML and one for optional Json metadata. The XML reader supports metadata as written by EPU and the optional Json matata reader can be used to read in data from other automated acquisition software of from manual acquisitions.

### Exporter

The remote browsing and transfer are implemented using the Qssh library. Only data from selected micrographs that have been fully processed (determined by the flags set by the scheduler) will be transferred. Multiple processes, which open multiple ssh channels for remote transfer, are used to transfer the data in order to use the full bandwidth of the network connection to the remote storage.

### Reporting

The report creation in CryoFLARE is based on the Lime report generator framework (http://limereport.ru). The reporting engine was extended to allow reading image files from disk and integrate them into a report. The model containing the micrograph metadata is used as data source for the report generator, allowing direct selection of any metadata parameter configured in settings for reporting.

## Discussion and Outlook

With the development of CryoFLARE we provide a tool for live assessment and optimization of data collection, which can help electron microscopy laboratories and facilities to increase the throughput of single-particle cryo-EM projects.

CryoFLARE is routinely used in our facility for data collection and already contributed to multiple cryo-EM structures (e.g. BK-VLP_scFv ^22^, nucleosome/DDB2 ^23^, BRISC ^24^) at FMI and Novartis. In addition, it has been used by ThermoFisher for demonstrating data acquisition capabilities of their instruments.

The software allows for prompter and more in-depth data analysis of acquired data, supporting the microscope operator in improving the acquisition strategy and reducing the amount of low-quality data recorded.

The data aggregation and reporting facilities support the users in the documentation of their EM experiments and help to improve the accountability for EM facilities.

The feedback mechanisms included with CryoFLARE are a first step towards more data driven decisions, where the data acquisition strategy is optimized continuously based on the results of the data analysis. The flexibility and extendibility of CryoFLARE together with its tight integration to data acquisition make it an easy-to-use test system for new acquisition and processing strategies, which will help to drive the evolution of the single-particle cryo-EM field.

## Software availability

CryoFLARE binaries for Linux and documentation can be found at www.cryoflare.org. The source code of CryoFLARE is licensed under the GNU General Public License (GPL) and is available from GitHub: www.github.com/fmi-basel/faim-cryoflare.git.

## Author contribution

AS conceived, developed and tested the software. SC helped test the software. AS, CG and NT wrote the manuscript.

## Acknowledgements

We would like to thank Alexandra Graff-Meyer, Christian Wiesmann, Honnappa Srinivas, Céline Be and Maryam Khoshouei from the FMI/Novartis electron microscopy competence center for fruitful discussions. We are also very grateful for Adriano Marra for help with computing and Bart van Knippenberg, Mazdak Radjainia and Fanis Grollios for their collaboration in interfacing CryoFLARE with EPU.

---

**Supplementary Figure 1:**
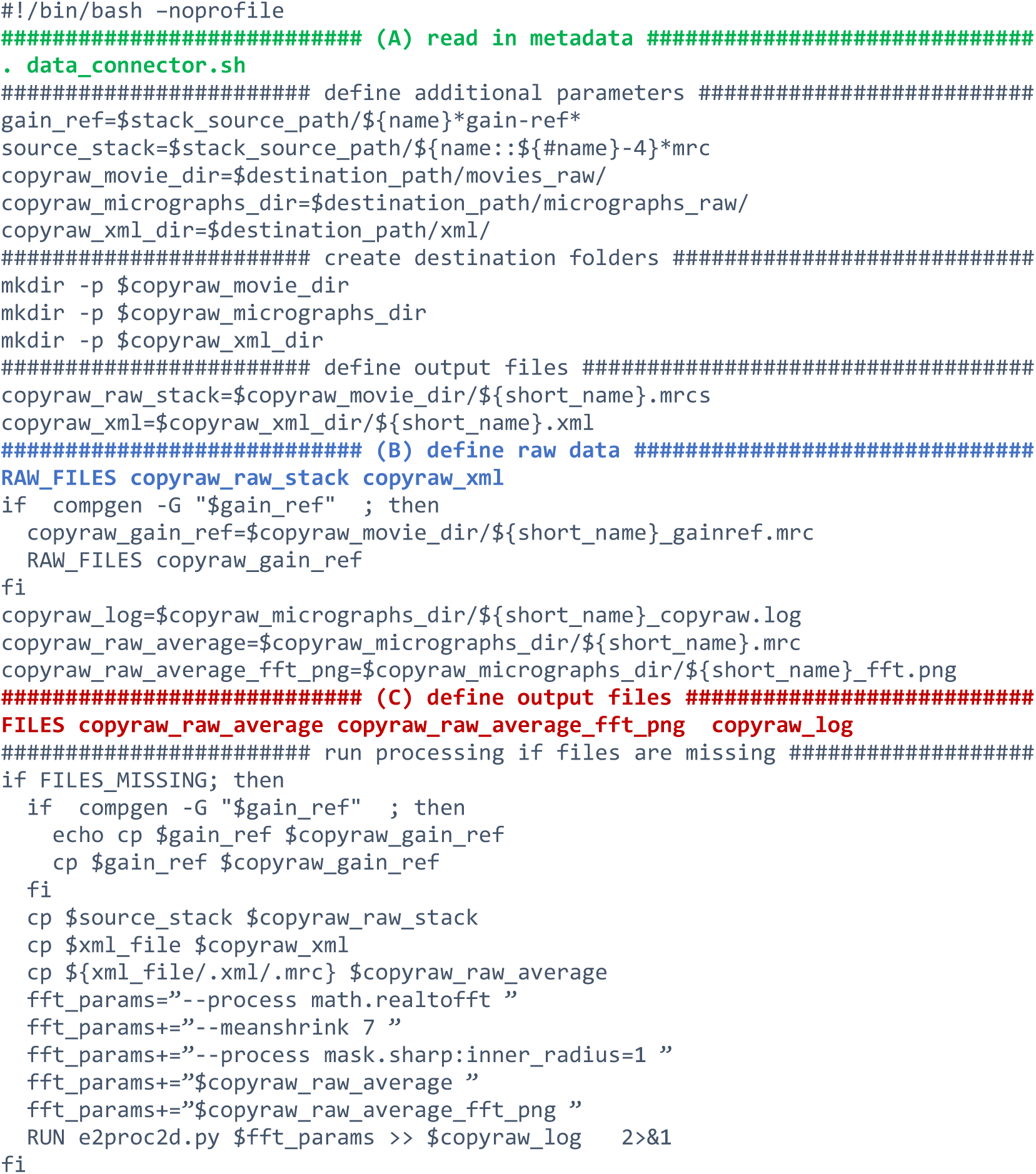
Raw data copy script. (A) adapter to read in metadata. (B) Function to define raw data output. (C) Function to define output files.

**Supplementary Figure 2:**
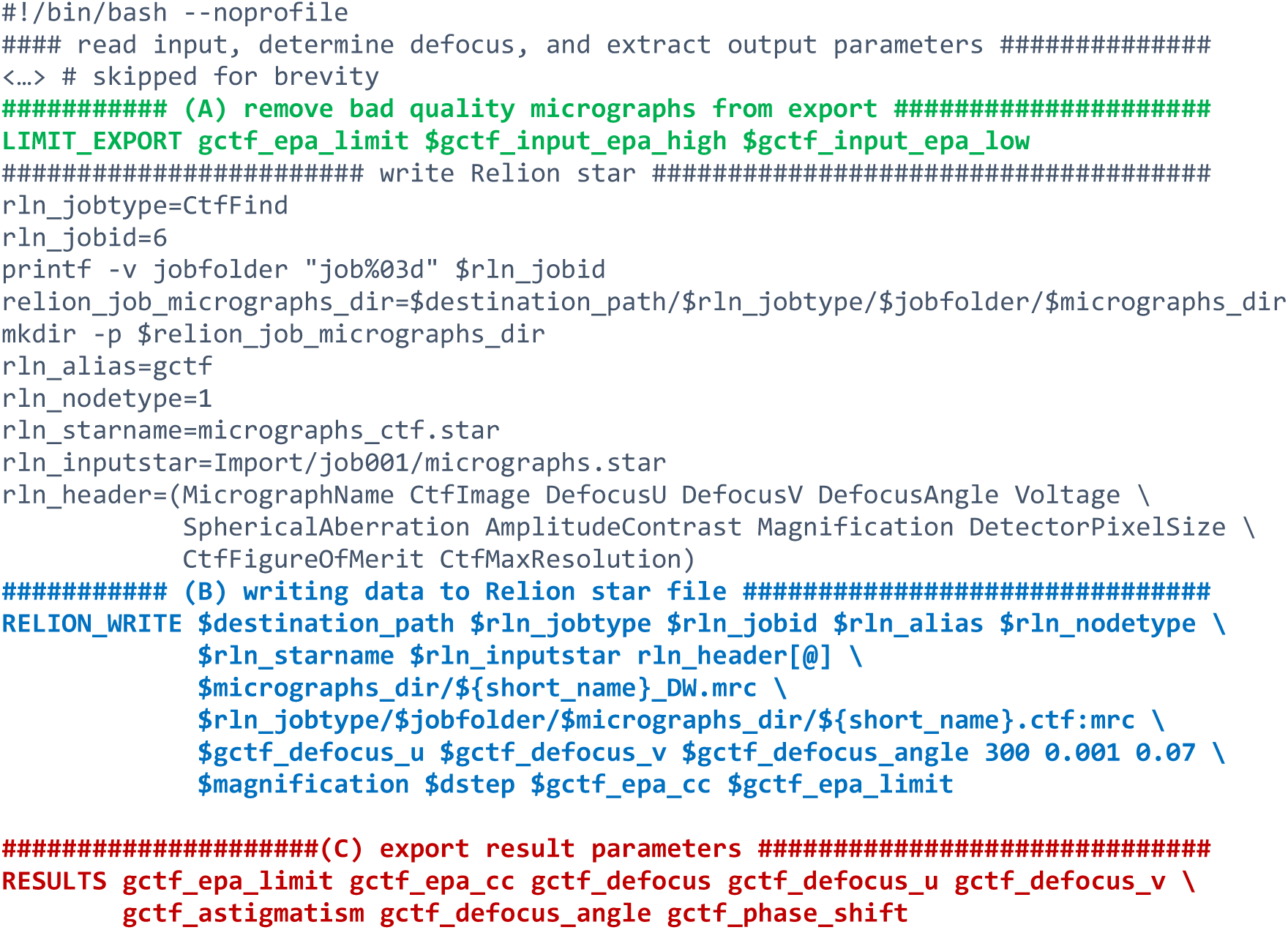
Data export part of CTF determination script. (A) data selection based on given limits. (B) Creation of Relion pipeline and star files and adding data to Relion star file. (C) Definition of result parameters.

